# Studies on a luminescent antimicrobial metal complex reveal a method to directly image pathogenic bacterial infection in a living organism using optical microscopy

**DOI:** 10.64898/2026.10.21.760849

**Authors:** Kirsty L Smitten, Hannah M Southam, Nick Van Hateren, Christa Walther, Jim A Thomas

**Affiliations:** Chemistry, School of Mathematics and Physical Science, University of Sheffield, Western Bank, Sheffield S3 7HF, UK; Molecular Biology and Biotechnology, School of Bioscience, The University of Sheffield, Western Bank, Sheffield S10 2TN, UK; Department of Biochemistry and Cell Biology, University of Vienna, Dr.-Bohr-Gasse 9, 1030 Wien, Austria

## Abstract

Methods to render *Galleria Mellonella* larvae transparent allowing three-dimensional viewing of larvae through orthodox light microscopy techniques are reported. To explore this model’s capability, we exploited the imaging properties of a metal complex antimicrobial to probe its therapeutic action against a pathogenic A*cetinobacter baumannii* infection. To illustrate the general utility of this method, we have also used the commercial fluorescent dye, Alexafluor-488, to image bacteria within live larvae.

## MATERIALS AND METHODS

### Injecting larvae

An overnight culture of *Acetinobacter baumannii*, AB184, was grown in MH-II media in an incubator at 37.5 °C. The culture was washed in phosphate buffered saline (PBS) (x3) and diluted to 10^5^ CFU/mL in dH_2_O. To stain the cells, AB184 were incubated with **1** (1.2 µM) for 30-minutes or Alexafluor-488 (0.5 µg/mL) for 5-minutes then washed in PBS (x3). Larvae were injected with the stained AB184 cells (10 µL) into their left pro-leg using a 1 mL Hamilton syringe. Syringes were cleaned in between each injection using the following method: water, ethanol, water. Once infected, larvae were left for 30-minutes before anaesthetising.

### Anaesthetizing larvae

It was found typical anesthetics caused darkening of the skin, melanisation, preventing optimal clearing. A self-made diethyl ether chamber proved the best anaesthetic method – Figure 1a main text. Cotton wool was soaked in diethyl ether and placed in a glass chamber. Larvae were incubated in the chamber for 20-30 minutes dependent on their size. Once anaesthetized larvae were fixed at room temperature using 4 % paraformaldehyde (PFA) in PBS. To fix, larvae were submerged into 1 mL of the PFA solution, leaving their head un-exposed.

**Figure 1.**
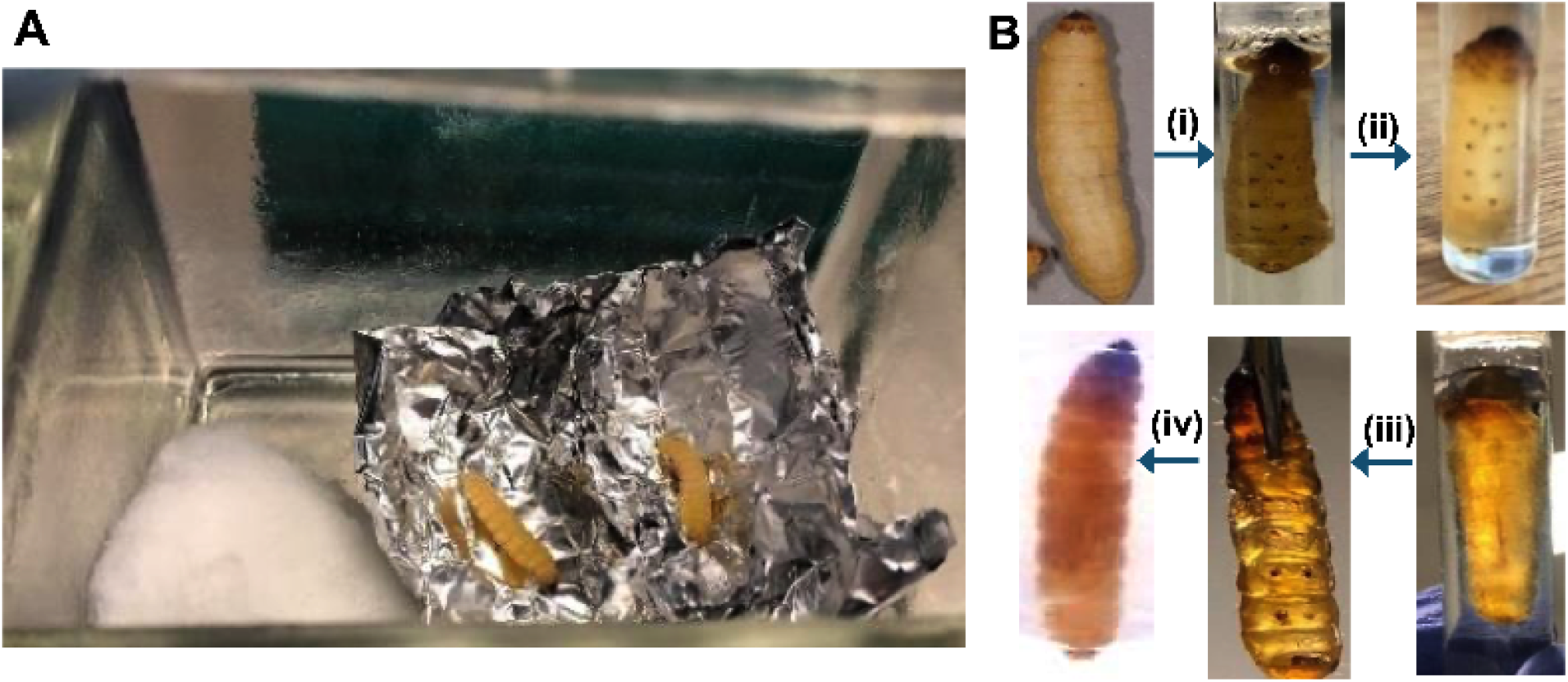
(**a**) Anesthetic chamber containing diethyl ether-soaked cotton wool. Larvae were incubated for 20-30 minutes at room temperature in the chamber. **(b)** Stages of clearing up to completely cleared larvae ready for imaging

### Clearing solutions

Solution one (CUBIC-1) was made up of urea (125 g) and Quadrol (156 g of 80 wt%) dissolved with hot stirring in 144 mL of dH_2_O. Once dissolved Triton X-100 (75 g) was added at room temperature. The solution was degassed with a vacuum desiccator (~0.1 MPa, ~30 min). CUBIC-1 was then mixed with dH_2_O (1:1), which is termed ½-CUBIC-1 in the clearing protocol. The reagent was stored at room temperature for a month, after this fluorescence signals can be affected during clearing. Solution two (CUBIC-2) was made by dissolving urea (12.5 g) and sucrose (25 g) in 7.5 mL of dH_2_O in a microwave. After complete dissolution the mixture was cooled to room temperature and triethanolamine (5 g) was added. The reagent was degassed with a vacuum desiccator (~0.1 MPa, ~30 min). CUBIC-2 was then mixed with PBS (1:1), which is termed ½-CUBIC-2 in the clearing protocol. The reagent was stored at room temperature for 2-weeks.

### Clearing protocol

**Day 1** - (i) Anaesthetised larvae were fixed using cold 1 mL 4 % (wt/vol) paraformaldehyde (PFA) in PBS (pH 7.4) for 1 hour at room temperature in an Eppendorf. PFA solution was used at pH 7.4 for an efficient clearing and to decrease larvae autofluorescence. Larvae were washed four times with cold PBS to remove fixant.

(ii) Fresh CUBIC reagents were used at the start of each clearing process. Larvae were incubated in 5 mL of ½-½-CUBIC-1 with rotation (~5 r.p.m) - Figure 1b manuscript – at 37.5 °C for 12 hours. A 5 mL Eppendorf was used over the 1 mL used in the previous step as clearing solutions can cause the tissue to swell.

**Day 2** - ½-CUBIC-1 was discarded and replaced with 5 mL CUBIC-1. The sample was rotated overnight at 37.5

**Day 3** - 5 mL CUBIC-1 was replaced, and larvae rotated at 37.5 °C. At this point significant clearing in the larvae’s skin is observed – Figure 1b manuscript. The CUBIC-1 reagent was then replaced every 2-days. Generally, depending on larvae size sufficient clearing for the next stage was observed within 10-14 days – Figure 1b manuscript.

**Day 14** - Larvae were then washed using 5 mL PBS/0.01 % (wt/vol) sodium azide three times, *via* rotation at room temperature for 3-hours each time. Larvae were then incubated in 5 mL ½-CUBIC-2 with rotation, for 24-hours at room temperature.

**Day 15** - The ½-CUBIC-2 was immediately replaced with 5 mL of CUBIC-2 and incubated with shaking, overnight at room temperature. A rocker was used over the rotation wheel in this step as CUBIC-2 forms bubbles when rotated. The following morning CUBIC-2 was replaced with fresh reagents and the larvae incubated with shaking for 24-hours.

**Day 17** - At this point smaller larvae were sufficiently clear; however, some larvae were incubated in CUBIC-2 for up to a further 7-days. Each day CUBIC-2 was replaced with fresh reagent.

### Imaging: larvae preparation

Cleared larvae were imaged in a 35 mm Cell Imaging Dish equipped with 145 µm glass thickness. Larvae were fixed into the dish using agarose, with pro-legs facing upwards. Samples must be prepared fresh on the day of imaging to prevent clearing reversal.

### Confocal Imaging

Larvae were imaged using an inverted Nikon A1 confocal system, using Nikon elements software. A CFI Plan Fluor 10x (NA 0.3) objective lens was used, for full larvae images a 5x objective would be optimal. Given the photochemical properties of 1 (excitation ʎ_max_: 450 nm, emission ʎ_max_: 600 nm) larvae were excited, and images were taken with the Red Fluorescent Protein (RFP) emission filter.

### Stereomicroscope Imaging

Larvae were imaged using a Nikon SMZ1500 stereomicroscope with a Prior Z-drive and transmitted and reflected illumination using Nikon elements software. A 1x objective was used. Microscope settings: NHS ester 488 (excitation ʎ_max_: 490 nm, emission ʎ_max_: 520 nm), **1** (excitation ʎ_max_: 490 m, emission ʎ_max_: 640 nm).

### Larvae disposal

Following imaging, larvae in agarose, over a number of days, reverted back to their opaque skin. Therefore, imaged larvae were disposed of, *via* freezing and disposal into a biological waste bin.

## INRODUCTION

The use of animal models has proved invaluable in research^1–3^ with invertebrate models being increasingly adopted for such studies,^4,5^ particularly as they are not subject to the ethical and practical limitations of a mammalian models.

Although *Drosphilla* have been used as models for bacterial infection,^6,7^ they require specialist housing and culture costs.

A promising alternative is the larvae of the wax moth *Galleria mellonella*. As larvae do not require husbandry costs, specialist equipment and culture costs are not incurred. Yet, *Galleria* provides a convenient model to study Gram-positive and Gram-negative bacterial infections.^8^ Although the larvae do not possess adaptive immunity, their innate immune system exhibits defenses similar to those of mammals.^9^ Furthermore, similar to human macrophages or neturophils, their hemocytes function as phagocytes, degrading bacteria in phago-lysomal compartments,^10,11^ Although infections and their treatment can be monitored within *Galleria*, one disadvantage they possess *versus* optically transparent models, such as *C elegans* and Zebrafish, is they are unsuited to studies involving light microscopy.

To develop a model of ESKAPE infections in a multicellular organism we identified a method to infect *Galleria* larvae with carbapenem-resistant *Acetinobacter baumannii*, CRAB.^12^ In studies on this model, we found that at lower concentrations of *A. baumannii*, AB184, larvae fight off infection through their innate immune system with the hemocyte cells ingesting/phagocytosing the bacteria. However, injection at or above the pathogenic concentration of 10^5^ CFU/mL the *Galleria* immune system no longer clears infection, resulting in significant larval death.We recently used this model to assess the therapeutic potential of previously identified^13,14^ antimicrobial complex, **1**.^12^ Here, we describe how this infection model can be extended to produce transparent *Galleria* that were used to visualize the progress of an infection by light microscopy.

## RESULTS AND DISCUSSION

An adaptation of the original CUBIC method,^15,16^ used in neurobiology research of brains from rodents, was developed to allow the larvae skin tissue to become transparent. This protocol, as with the original, involves a two-step incubation of fixed tissue in two separate aqueous solutions. However, the protocol for rodent tissue involves incubation for one-two weeks, larvae clearing takes 2-3 weeks depending on the size of the larvae. In this protocol, – see methods for detail – individual *Galleria*.larvae were cleared – Fig 1.

To illustrate the facility of this process, using the luminescence of complex **1**, we first investigated the uptake of -and immune response to-infecting bacteria within cleared *Galleria*. In these studies, the CRAB bacterial strain was first labelled with **1**, then injected into the larvae at a pathogenic concentration (10^6^ CFU/mL), which were then incubated for 30-minutes. Following anaesthetization with diethyl ether, the larvae were incubated in PFA (4 %) for a further 30 minutes and then cleared using the adapted CUBIC protocol. This enabled sites of infection and effects on larvae hemolymph cells to be visualized-Fig 2.

**Figure 2:**
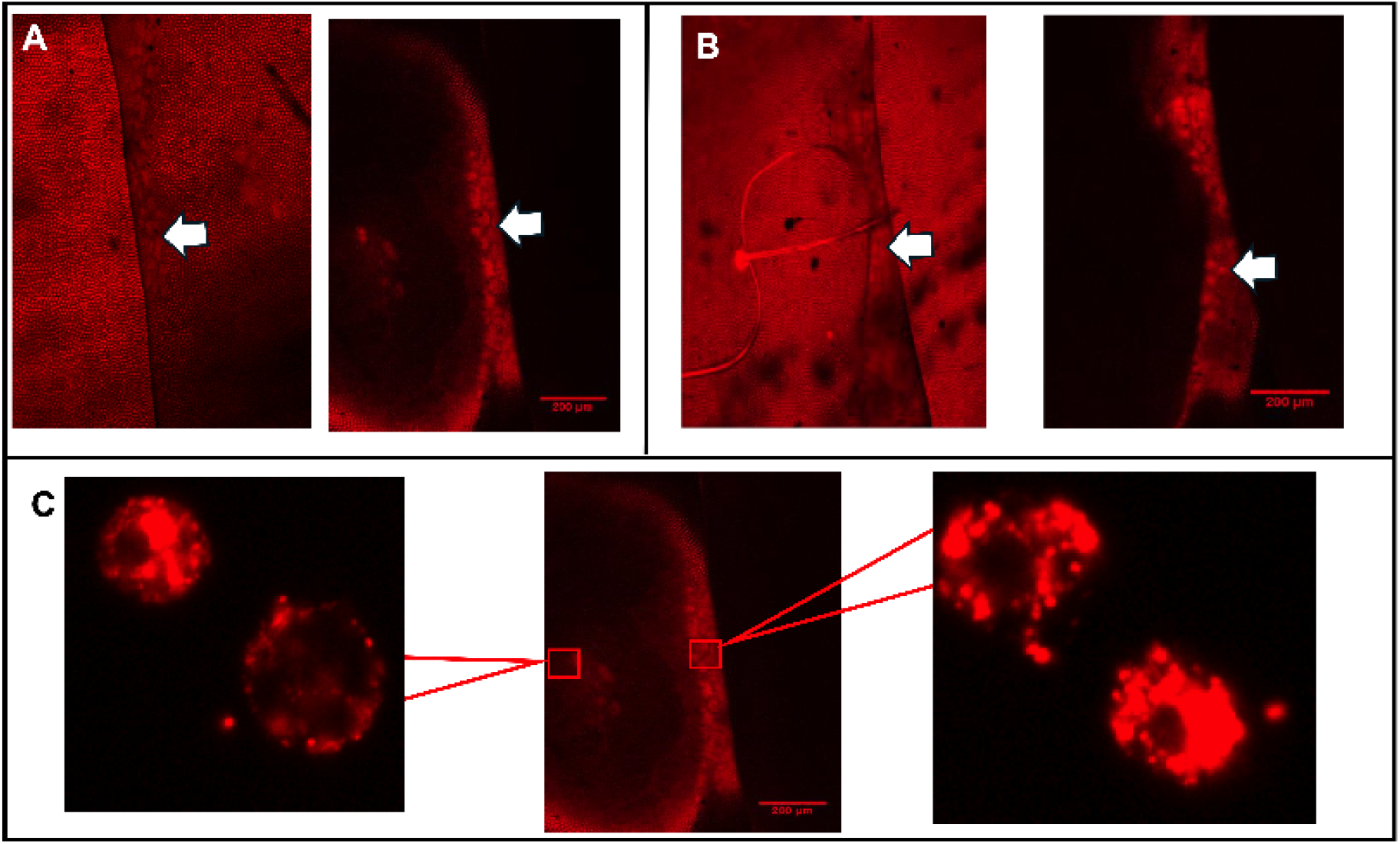
Bacteria stained with **1** within *Galleria* larvae; *A. baumannii* cells are highlighted by arrows. **A**. Images of head region **B**. Analogous images of tail region. **C**. Images showing initiation of an immune response in *Galleria* hemolymph cells.

To investigate whether this protocol is more generally applicable, Alexafluor-488 (3A) was then used to stain bacteria and compared to images generated with **1**. To confirm clearing could be used to visualize *whole* larvae, images were taken on a fluorescent stereomicroscope – Fig 3. Again, bacteria were seen to localize within the central hemolymph running through the larvae

**Figure 3.**
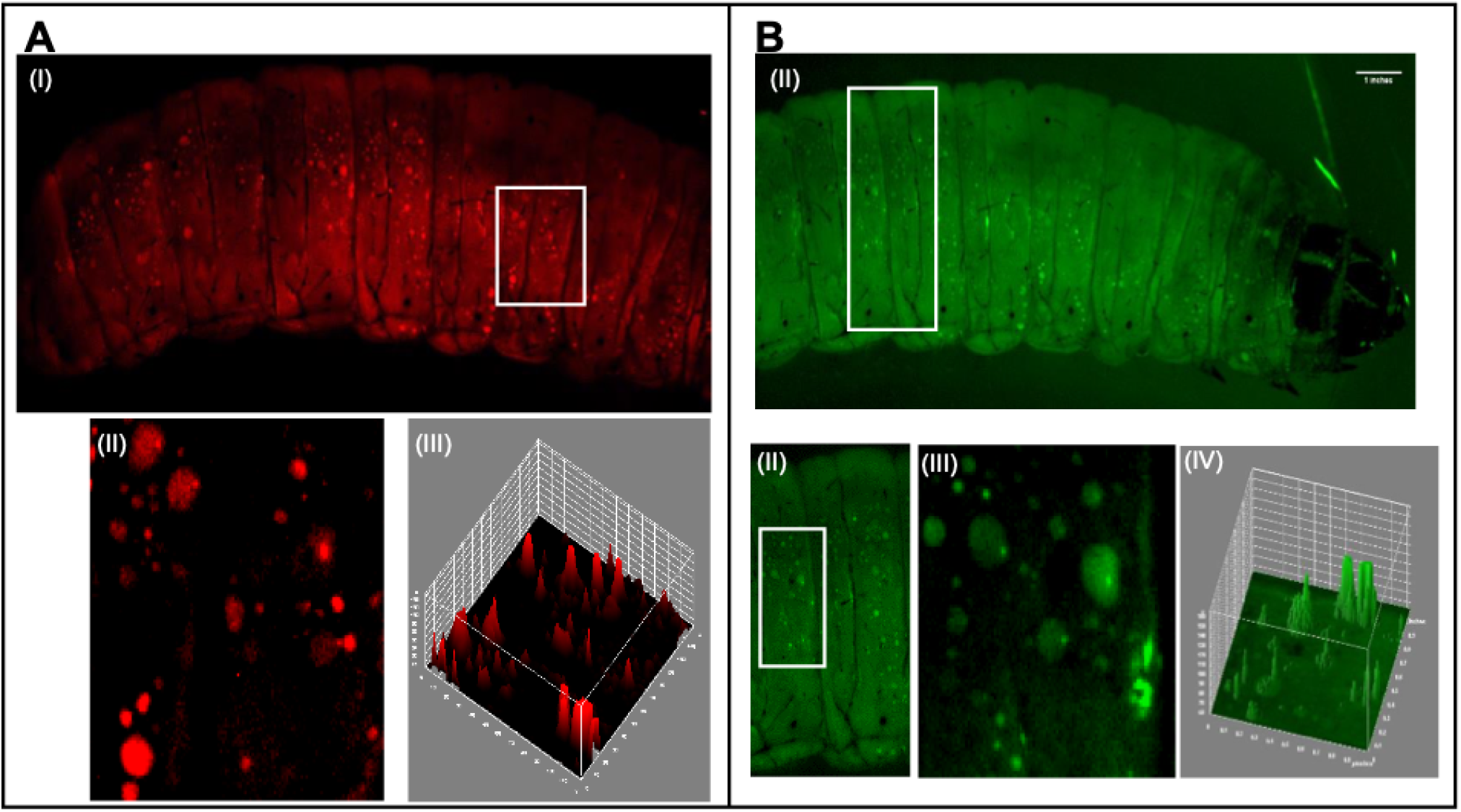
Whole larvae imaging. **A**. (I) Larva stained with **1** (II) Expansion of white box in(I) to show bacteria within hemocytes. (III) 3D surface plot show peak emission intensity in (II). **B**. (I) Larva stained with Alexafluor 488 (0.5 mg/mL). (II) and (III) Expansion of white box in previous panel revealin bacteria within hemocytes. (IV) Accompanying 3D surface plot for (III).

Interestingly, within a few hours of CUBIC treatment, its effect is reversed and eventually larvae became fully melanized, indicating that they are still metabolically active even *after* CUBIC treatment. To avoid this effect, it is recommended that larvae are imaged ~20-minutes after clearing.

## CONCLUSIONS

Summing up, by adapting CUBIC clearing methods, a protocol that renders *Galleria* larvae optically transparent is reported. To illustrate the applicability of this method, it was used to investigate a CRAB infection and its treatment within larvae from a cellular to whole organism scale. As *Galleria* are increasingly utilized to investigate a range of pharmacological issues, from estimating *in vivo* toxicity, through assessing microbial pathogenicity, to the identification of new therapeutics, the application of optical microscopy/nanoscopy in this model offers potential to carry out an even wider range of studies with a greater depth of understanding. Finally, as the whole organism and sites of infection can be simultaneously directly imaged at a range of scales, this approach provides an advantage over other methods of *in vivo* monitoring of pathogenic infections and putative treatments.

## Acknowledgements

We are grateful to the BBSRC White Rose Structural Biology DTP (BB/M011151/1; studentship to KS). HMS acknowledges BBSRC grant (BB/M022579/1). The Wolfson Light Microscopy Facility was funded by MRC grant MK/K0157531/1.

## Author Contributions

KLS: conceptualization, investigation, methodology, validation, writing – original draft preparation; HMS: conceptualization, investigation, methodology,resources, writing – original draft preparation; NVH: investigation, resources, validation, visualization; CW: investigation, resources, validation, visualization, supervision; JAT: conceptualization, funding acquisition, project administration, supervision, visualization, writing – original draft preparation, writing – review and editing.

## Notes

### Competing Interest Statement

The authors have declared no competing interest.

